# Reduced triacylglycerols and lipid droplets are associated with resilience to Alzheimer’s disease

**DOI:** 10.1101/2025.05.08.652735

**Authors:** Daan van der Vliet, Luuk E. de Vries, Xinyu Di, Brechtje de Jong, Marc T.C. van der Meij, Marielle van der Peet, Amy C. Harms, Thomas Hankemeier, Dick F. Swaab, Inge Huitinga, Joost Verhaagen, Mario van der Stelt

**Affiliations:** Department of Molecular Physiology, Leiden University & Oncode Institute, Leiden 2333 CC, Netherlands; Department of Neuroimmunology, Netherlands Institute for Neuroscience, Institute of the Royal Netherlands Academy of Arts and Sciences, Meibergdreef 47, 1105 BA, Amsterdam, The Netherlands; Department of Neuroregeneration, Netherlands Institute for Neuroscience, Royal Netherlands Academy of Arts and Sciences, Meibergdreef 47, 1105 BA Amsterdam, The Netherlands; Metabolomics and Analytics Centre, Leiden Academic Centre for Drug Research, Leiden University, Leiden 2333 CC, Netherlands; Department of Pathology, Amsterdam Neuroscience, Amsterdam UMC - Location VUmc, Amsterdam, The Netherlands; Department of Neuropsychiatric Disorders, Netherlands Institute for Neuroscience, Institute of the Royal Netherlands Academy of Arts and Sciences, Meibergdreef 47, 1105 BA Amsterdam, The Netherlands

**Author notes:** these authors contributed equally (Daan van der Vliet, Luuk E. de Vries).

**Keywords:** Lipid metabolism, Alzheimer’s Disease, Resilience, lipid droplets, neuroinflammation

## Abstract

**Introduction:** While it has become clear that alterations in lipid metabolism are associated with AD, it is unclear how they contribute to both cognitive decline and the pathophysiology of AD.

**Methods:** We performed lipidomics and activity-based protein profiling in the frontal cortex of control, AD and resilient donors, i.e. individuals with AD pathology without cognitive decline. Subsequently we integrated these data using multi-omics factor analysis and correlated the multi-omics profiles to disease and clinical parameters.

**Results:** The most pronounced alterations in lipids were in the ω6-derived oxylipins, which were particularly increased in the AD patients. Both triacylglycerols (TAGs) and lipid droplets were more abundant in the AD donors compared to the resilient donors. Enzyme activities showed a similar direction in the AD and resilient donors, including decreased activity of ABHD6. Multi-omics factor analysis showed that increased oxylipins, loss of inhibitory cells, synaptic genes and genes related to the inflammatory response were associated with Aβ plaque load in both AD and resilient donors.

**Conclusion:** Our multi-omics data show a response associated with Aβ load shared among AD and resilient donors and, for the first time, reduced lipid droplets in resilient donors.

## Introduction

Alzheimer’s Disease (AD) is the most common form of dementia, severely affecting memory and other cognitive functions. Effective therapies for the prevention of cognitive decline are currently lacking, and thus there is an urgent need for new therapeutic targets for the development of novel therapies. Recently, the FDA approved several monoclonal antibodies targeting amyloid-β (Aβ) plaques. However, the clinical efficacy of these antibodies is limited and they cause serious side-effects, like oedema and frequent haemorrhages [1]. Thus, it is likely that removing Aβ in AD patients alone is not sufficient to halt cognitive decline. This is supported by the fact that some individuals are cognitively intact despite having considerable amounts of AD neuropathology, a phenomenon also known as resilience [2–5]. It is thought that these resilient individuals have a different cellular response to pathology, as opposed to AD patients, in which these cellular responses may initiate a cascade leading to cognitive decline. Studies further investigating the possible responses to AD pathology in resilient individuals and AD patients are of potential therapeutic interest.

Recently, our understanding of the cellular and molecular responses to AD pathology in both AD patients and resilient individuals has improved by studying alterations in gene expression and their proteome. Numerous studies have demonstrated a shift towards more reactive glial cells as a reaction to AD pathology [6–9], synaptic vulnerability to oligomeric p-tau and Aβ species and mitochondrial dysfunction. Interestingly, such changes are often absent or less pronounced in resilient individuals, which could at least partly explain why these individuals remain cognitively intact [10–13]. While these changes are apparent at the gene expression or protein levels, changes in other biomolecules, such as lipids, are not well studied.

Changes in lipid metabolism have been implicated in many processes related to aging and AD. Lipids play crucial roles in neuronal function, energy regulation, myelination and neuroinflammation, processes which are often altered in AD [14, 15]. Interestingly, large-scale genome wide association studies in AD patients have identified multiple risk loci related to cholesterol and lipid metabolism, including Apolipoprotein E (ApoE) [14]. More recently, lipid-droplet-accumulating microglia surrounding plaques have been described in animal models for AD and in AD patients [16–18], which were associated with increased levels of neuroinflammation [17].

Besides playing roles in structural membrane formation and energy demands, lipids are also increasingly recognized as signalling molecules. Lipids like endocannabinoids, lysophosphatidic acids, prostaglandins and sphingosine-1-phosphate, contribute to synaptic plasticity, memory formation, oligodendrogenesis, bouton formation and excitability [19–24]. Furthermore, lipids are involved in neuroinflammatory responses through arachidonic acid (AA) derived neuroinflammatory prostaglandins and other oxylipins [22]. Various steps in the cascade for AA production and subsequent oxidation to oxylipins have previously been shown to be altered in AD and AD mouse models [21, 25–28]. However, it remains unclear if lipid signalling contributes to or is altered by the pathophysiology of AD, and how this relates to cognition.

Here, we have applied lipidomics and activity-based protein profiling (ABPP), a methodology to identify active lipid metabolizing enzymes, on grey matter tissue from the frontal cortex of a group of well-characterized AD patients, resilient individuals and matched control donors to investigate alterations in lipid metabolism. Furthermore, we integrated both datasets with a previously published bulk RNA sequencing dataset to perform a multi-omics analysis, linking cell types and synaptic processes to changes in lipids. Finally, we confirmed the findings of the lipidomic and ABPP analyses using specialized fluorescent probes, western blot and immunohistochemistry.

## Materials & methods

### Donors & Tissue

Human brain tissue was obtained from the Netherlands Brain Bank (NBB). Informed consent for use of brain material and clinical data for research purposes and brain autopsies was obtained by the NBB according to international ethical guidelines. Autopsy procedures were approved by the Medical Ethic Committee of the AUMC, Amsterdam, the Netherlands. Neuropathological assessments were done according to standardized protocols [29]. Similar donors were selected as described previously [30]. In brief, donors were selected based on the amount of AD neuropathology and cognition (clinical dementia rating scale (CDR) or global determination scale (GDS)): Demented AD patients (CDR 3, Braak 4-6, Thal ≥ 3), resilient donors (CDR ≤0.5, Braak 3-5, Thal ≥ 3) and age-matched controls (CDR ≤0.5, Braak 1-2, Thal ≤ 2). Cases with signs of neurological or psychiatric diseases other than those associated with AD were excluded. Donors with severe comorbid pathology (e.g. cortical Lewy bodies, TPD-43, hippocampal sclerosis) were excluded. Donors were matched as closely as possible for sex, age, pH, post-mortem delay (PMD) and ApoE genotype (Table 1).

**Table 1:** Donor characteristics.

|  | Control | AD | Resilient | p-val. |
| --- | --- | --- | --- | --- |
| N | 12 | 12 | 12 |  |
| Sex | 6M/6F | 6M/6F | 6M/6F | >0.999 |
| Age | 82.2 ± 10.7 | 82.2 ± 9.9 | 88.6 ± 8.4 | 0.188 |
| PMD | 6.0 ± 1.8 | 5.2 ± 1.0 | 6.5 ± 1.6 | 0.119 |
| pH | 6.6 ± 0.3 | 6.5 ± 0.2 | 6.4 ± 0.2 | 0.061 |
| Braak | 1.3 ± 0.7 | 5.5 ± 0.8 | 4.1 ± 0.9 | <0.001 |
| Aβ plaque score (A) | 0.7 ± 0.5 | 2.9 ± 0.3 | 2.4 ± 0.5 | <0.001 |
| Neuritic plaque score (C) | 0.1 ± 0.3 | 2.8 ± 0.6 | 1.4 ± 0.8 | <0.001 |
| ApoE4+/- | 3+/9- | 7+/5- | 6+/6- | 0.441 |

Abbreviations: AD, Alzheimer's disease; PMD, post-mortem delay; ApoE4+/-, presence or absence of Apolipoprotein E 4 allele; p-val, p-value.

### Lipidomics sample preparation

Snap frozen human brain tissue from the superior frontal gyrus (SFG) was cut on a cryostat. For each donor, between 15 and 25 mg of tissue was isolated by sectioning ∼10 sections (50 µm). Grey matter was dissected out inside the cryostat using pre-chilled scalpels, collected in pre-chilled tubes and immediately put on dry ice. The tissue was homogenized by mechanical lysis with 0.5 mm glass beads in ice-cold lysis buffer (20 mM HEPES, 1 mM MgCl_2_, 1 U/mL benzonase). 20 µL of lysis buffer/mg tissue was used. The protein concentration was measured using a Bradford assay (Bio-Rad), the lysates were diluted in lysis buffer to a concentration of 1 mg/mL and aliquots of the lysates were snap frozen in liquid nitrogen and stored at-80.

Purchased or synthesized standards/internal standards (IS) were dissolved in methanol, ethanol, chloroform or ACN at different stock concentrations. These stock solutions were further diluted and mixed to make the standard stock solutions and IS stock solutions. The lipids IS mix contained deuterated version of ceramides, (lyso)phospholipids, diacylglycerols, triacylglycerols and cholesterol esters. The signaling IS mix contained deuterated version of oxylipins, endocannabinoids (eCBs), free fatty acids and bile acids. The signalling calibration standards contained oxylipins, eCBs, free fatty acids and bile acids.

The aliquots of sample lysates (∼1 mg protein/mL, 78 μL) were thawed on ice in 1.5 mL Eppendorf tubes. To each sample 10 μL IS work solution was added. Calibration samples were prepared by spiking 10 μL of each calibration standards into 78 μL of water. Extraction was performed by 100 μL extraction buffer (0.2mM Ammonium formate) and 1000 μL extractant (BuOH:EtOAc, 50:50, v/v). Samples were then mixed in a Next Advance Bullet Blender (5 min, room temperature) followed by centrifugation (16,000 x g, 10 min, 4°C). 900 μL of the organic layer was transferred into clean safelock low-binding tubes and concentrated in a SpeedVac vacuum concentrator (45 ⁰C, ThermoFisher), followed by adding 60 µL of reconstitution solution (MeOH:ACN, 30:70, v/v) and agitating for 15 min. The reconstituted samples were centrifuged (16,000 g, 10 min, 4°C) and 50 µL were transferred into autosampler vials with inserts. Samples were kept at-80 °C till LC-MS analysis.

### Lipidomics LC-MS/MS measurements

Samples were randomized and run in one batch on three LC-MS/MS platforms. Each batch included QC samples and blank samples. QC samples are used to assess data quality. Each batch included study QC samples. Method blanks (proc blanks) are used to check for background signal. These samples are composed of analyte-free matrix and have gone through all the steps of the sample preparation procedure with the reagents only.

The signalling lipids platform covers 196 metabolites, including prostaglandins and other oxylipins, N-acylethanolamines, monoacylglycerols and free fatty acids. Bile acids are also included in this platform but were not detected. Reference standards were used for each analyte for peak identification, 47 deuterated internal standards were used for the correction of variations from sample preparation and LC-MS runs. A QTRAP 7500 (AB Sciex, Concord, ON, Canada) coupled to an Exion LC AD (AB Sciex, Concord, ON, Canada). MS/MS experiments were done with a Turbo V source (AB Sciex, Concord, ON, Canada) operated with ESI probe. An Acquity UPLC BEH C18 column (Waters) was used to measure the samples. The three-pump LC system consisted of mobile phase A (MPA, H2O with 0.1 % acetic acid), mobile phase B (MPB, 90 % CH3CN /10 % MeOH with 0.1 % acetic acid) and mobile phase C (MPC, IPA with 0.1 % acetic acid). The injection volume was 5 μL sample stacked with 10 μL of mobile phase A. The flow rate was 0.7 mL/min, and each run takes 16 min. The gradient started at 20 % MPB and 1% MPC. The MPB progressed from 20% to 85% between 0.75 min and 14 min while the MPC ascended from 1 % to 15 % between 11 min to 14 min after which conditions were kept for 0.3 min and then the column was re-equilibrated at initial conditions until 16 min. An electrospray ionization source (ESI) was used with parameters: interface temperature 600 °C, curtain gas 45 psi, CAD gas 9 psi, gas 1 and gas2 both 65 psi. The mass spectrometer operated in polarity switching mode and all analytes were monitored in dMRM mode. Data was acquired using Sciex OS Software V2.0.0.45330 (AB Sciex).

The phospholipids platform covers 1320 nonpolar lipids targets, based on a previously described method [31]. The samples were measured using 3 separate acquisition methods with same LC-MS conditions but different MRM transitions. Acquisition method 1 measurement contains classes of phosphatidylcholine, phosphatidylinositol, phosphatidylserine, phosphatidylglycerol, and bis(monoacylglycerol)phosphates. Acquisition method 2 measurement contains sphingomyelins, hexosylceramides, lactosylceramides and phosphatidylethanolamines. The acquisition method 3 measurement contains triacylglycerols. The identification of metabolites in this platform are based on the dimensions of specific MS/MS transitions and retention time (RT), Using HILIC columns, the lipids from the same class elute in a narrow RT window, while different lipid classes elute at different RTs. For each class, one or more deuterated internal standards were used to check the RTs and variations from sample preparation and LC-MS runs.

A QTRAP 6500+ (AB Sciex, Concord, ON, Canada) was coupled to an Exion LC AD (AB Sciex, Concord, ON, Canada). MS/MS experiments were done with a Turbo V source (AB Sciex, Concord, ON, Canada) operated with ESI probe. A Phenomenex Luna® amino column (100 mm × 2 mm, 3 μm) was used for separation. The mobile phase A was 1 mM ammonium acetate in chloroform: acetonitrile (1:9), while mobile phase B was 1 mM ammonium acetate in acetonitrile: water (1:1). The gradient started at 0 % MPB. The MPB progressed from 0 % to 50 % between 2.1 min and 11 min, from 50 % to 70 % between 11 min and 11.5 min, kept at 70 % for 1 min, and then the column was re-equilibrated at initial conditions from 12.6 min to 14 min. The injection volume was 2 μL for all the three injections. The column temperature was kept at 35°C. The injector needle was washed with isopropanol:water:dichloromethane (94:5:1, v:v:v) after each injection. Assigned MRM peaks from the acquired data were integrated using SCIEX OS (version 2.1.6) Software and signals were corrected using proper internal standards.

The RP MS/MS-based lipids platform covers 186 lipids, including ceramides, diglycerides, and cholesterol esters. The identification of metabolites in this platform are based on the dimensions of specific MS/MS transitions and retention time (RT), Using reversed phase columns, the lipids from the same class elute at RTs that can fits into linear regression models involving carbon number and double bond number (RT mapping). For each class, one or more deuterated internal standards were used to check the RTs and variations from sample preparation and LC-MS runs.

A QTRAP 7500 (AB Sciex, Concord, ON, Canada) coupled to an Exion LC AD (AB Sciex, Concord, ON, Canada). MS/MS experiments were done with a Turbo V source (AB Sciex, Concord, ON, Canada) operated with ESI probe. An Acquity UPLC BEH C8 column (Waters) was used to measure the samples. The mobile phase was consisted of 2 mM HCOONH_4_, 10 mM formic acid in water (A), ACN (B), IPA (C). The gradient was the following: starting conditions 10% B and 10% C; increase of B from 10% to 40% between 1 min and 2 min; maintaining B at 40% and C at 10% between 2 min and 7 min; increase of C from 10% to 45% between 7 min and 8 min; maintaining B at 40% and C at 45% between 8 min and 10 min; returning to initial conditions at 10.5 min and re-equilibration for 1.5 min. The triple quadrupole mass spectrometer operated in polarity switching mode and all analytes were monitored in dMRM mode. Data was acquired using Sciex OS Software V2.0.0.45330 (AB Sciex).

Assigned MRM peaks from the acquired data from all platforms were integrated using SCIEX OS (version 2.1.6) Software and signals were corrected using proper internal standards. Blank effects (BE) for each analyte were checked by comparing proc blank samples to quality control (QC) samples. The threshold for blank effects was 40%. The precision and reproducibility of the analytical process were checked using the relative standard deviations (RSDs) of the QCs. The threshold for QC-RSD was 30%. Any lipid with a blank > 40% and RSD > 30% was removed from the dataset.

## Data analysis & statistics

All downstream analysis was performed in R v4.4.1 using Rstudio. Basic processing and data handling were performed using the tidyverse framework (v2.0.0), magrittr (v2.0.3), statistics were calculated using rstatix (v0.7.2) and ggpubr (v0.6.0), and all plotting was done using ggplot2 (v3.5.1), ComplexHeatmap (v2.2.0), cowplot (v1.1.3) and ggrepel (v0.9.5).

### Processing of lipidomics data

Lipid levels are expressed as response ratio’s (integrated MRM peaks). The data were further normalized by the sum of all lipids and subsequently log2 transformed and centered. Missing values were imputed by sampling randomly from a distribution with a mean corresponding to the lowest value detected for a given lipid, and a standard deviation of 1/3 of that lipid. This is based on the assumption that if a lipid was not detected, it likely fell below the detection threshold, meaning that the concentration is relatively low.

### Activity-based protein profiling using chemical proteomics

The chemical proteomics workflow was based on the previously reported procedures [32]. Lysates were prepared as described above for lipidomics.

Lysates (100 µL, 1 mg/mL protein) were thawed on ice. 1% SDS was added to the negative controls and heated to 95 ⁰C for 5 minutes. Probe cocktail (0.5 mM FP-biotin and 0.5 mM THL-biotin) was added to the samples and negative controls (ratio 1:50) and incubated for 30 min at 37 ⁰C (800 rpm). The reaction was quenched by chloroform/methanol precipitation. 250 μL H_2_O, 465 μL MeOH, 120 μL CHCl3 and, 105 μL H_2_O was added to all samples and after each addition the samples were vigorously vortexed. The samples were centrifuged (1,500 g, 5 min, RT) and the upper aqueous phase was removed. The pellet and the lower chloroform phase were resuspended in MeOH (500 μL), and the pellet was resuspended by sonicating (10 % amplitude, 2×10 s). The methanol was removed after centrifugation (20,000 g, 5 min, RT) and the pellet redissolved in 250 μL PBS (0.5% SDS, 5 mM DTT). The samples were sonicated (10% amplitude, 2×10s) and kept at 65 ⁰C for 15 minutes while shaking (800 rpm). After cooling down to rt, 15 μL IAA (0.5 M iodoacetamide in Milli-Q) was added and samples were incubated for 30 min in the dark. The reaction was quenched by adding 5 μL DTT (1 mM DTT), vortex and centrifugation (20,000 g, 2 min).

From this point on, samples were handled in a flow cabinet, and hairnets and gloves were used to prevent sample contamination. 10 μL slurry (suspension of 50% high capacity streptavidin beads, Thermofisher 20361) and 30 μL slurry (suspension of 50% control agarose beads, Thermofisher 26150) were used for each sample. The beads were washed twice in 6 mL PBS (0.5% SDS) by vortexing and centrifuging (3,000 g, 2 min). And washed once with 6 mL PBS by vortexing and spinning down (3,000 g, 2 min). The beads were resuspended in PBS and divided in 1.5 mL tubes with 250 μL for each sample. 250 μL of the samples were transferred to the 1.5 mL tubes with beads. The beads were agitated by overhead turning (2 hours, RT). Afterwards, the samples were centrifuged (3,000 g, 2 min) and the supernatant discarded by pouring out. The beads were washed 4x with PBS with 0.5% SDS, vortexing, centrifuging (3,000 g, 2 min) and discarding supernatant. This step was repeated 5x with PBS to remove SDS. The beads were washed in digestion buffer (100 mM Tris pH8, 100 mM NaCl, 1 mM CaCl2 and 2% (v/v) ACN), centrifuged (3,000 g, 2 min) and supernatant removed by pipetting. The beads were resuspended in 100 μL digestion buffer containing 0.25 μg trypsin (Promega), and protein was digested overnight at 37 ⁰C while vigorously shaking (1000 rpm). To stop the digestion, 100 μL 10% formic acid (FA) in Milli-Q was added and centrifuged (3,000 g, 2 min). The samples were prepared by stage-tip column over an Oasis plate (Waters), which was conditioned by washing sequentially with MeOH, 3:2 ACN/H_2_O with 0.5% FA (solution B) and finally with H_2_O with 0.5% FA (solution A). The peptides were loaded onto the stagetips, washed with solution A and then eluted into clean safelock low-binding tubes (Eppendorf). The collected samples were dried in a speedvac (45 ⁰C, 2-3 h) and stored at-80 °C ⁰C until further use.

The peptides were redissolved in 30 μL LC/MS solution (Milli-Q/ACN/FA in 97:3:0.1 containing 10 fmol/μL yeast enolase (Waters, product no. 186002325; UniProt P00924)), by vortexing and spinning down briefly. peptides were then analyzed on a QExactive LC-MS/MS (Thermo Fisher). All gradients and solutions were according to previously published procedures [32]. Raw spectral data was analyzed by MaxQuant software (v2.0.1.0) [33, 34]. The ‘proteingroups.txt’ output file from MaxQuant was imported in R. Identified proteins were filtered for potential contaminants identified by MaxQuant, and additional criteria for the proteins were that 1) a protein was identified with 2 independent unique peptides, 2) The ratio of protein raw intensity of native, heat inactivated lysate was at least 2.0 in at least 5 out of 10 QC pairs. 3) The protein is annotated as a serine hydrolase, or has an annotation in Uniprot as “charge-relay system” or “nucleophile” as a catalytic residue and 4) The protein had an LFQ value for more than 60% of the samples in at least one group. 82 enzymes met these criteria and were further processed. Missing values were not imputed.

### PCA

Principal component analysis was performed using the ‘prcomp’ function in base R or the PCAtools package (v2.10.0) with scale set to FALSE and center set to TRUE. Number PCs to keep was determined using the ‘getelbow’ function and correlation of PCs to metadata was done with the ‘eigencorplot’ function of PCAtools, using Spearman correlations, with BH multiple testing correction.

### PLS-DA

Partial least squares – discriminant analysis (PLS-DA) was performed using the ‘ropls’ (v1.30.0) package in R, using pathology (control versus AD+resilient) as a response and Cognition (AD versus control + resilient) as a response. The full imputed lipidomics dataset was the data input, further settings were predI = 1 and scale=center. The obtained 2 dimensions were plotted against each other and data points were coloured according to group. Loadings were extracted from the PLS-DA and plotted on their respective axis using the ComplexHeatmap package [35].

### Differentially altered lipids and proteins

To calculate differentially abundant lipids or proteins from ABPP we used Wilcoxon rank sum tests. This test was chosen because >50% of lipids showed a non-normal distribution as judged by Shapiro tests. Multiple testing correction was performed using the Benjamini-Hochberg (BH) method accepting a false discovery rate of 25%. In volcanoplots, the raw Wilcoxon p-value is plotted, but coloured according to whether it is still significantly different after BH correction.

ROAST was used to calculate enrichment of lipid sets within the Limma framework (v.3.60.4) [36, 37]. The linear model was constructed using a simple linear model design y = ∼0 + group, where group corresponds to control, AD or resilient, and fed into the roast() function, with indexes referring to specific lipid classes, nrot was set to 10^4^ and set.statistic was set to “mean”.

### Gel-based activity-based protein profiling (ABPP) & western blot

Lysates were prepared as described above. 0.5 µL of a 40x concentrated stock of fluorescent probe in DMSO was added to 19.5 µL of lysate. The used probe was LEI-612-BDP-TMR (200 nM, 30 min 37 ⁰C). This probe was synthesized in-house according to published procedures [38, 39]. Full characterization of LEI-612-BDP-TMR will be described elsewhere. After incubation, the lysate was denatured through the addition of 7.5 µL 4xLaemmli buffer (final concentrations 60 mM Tris (pH 6.8), 2% (w/v) SDS, 10% (v/v) glycerol, 1.25% (v/v) β-mercaptoethanol, 0.01% (v/v) bromophenol blue) and 15 minutes incubation at RT. Proteins were resolved on an SDS-PAGE (10% acrylamide, 29:1 acrylamide:bisacrylamide, BioRad) for ∼75 minutes at 180V. Subsequently, gels were scanned for in-gel fluorescence on a chemidoc MP (Bio-Rad) on settings Cy2 (emission 532/50, 120s exposure), Cy3 (emission 600/50, 120s exposure) or Cy5 (emission 700/50, 20s exposure).

The SDS-PAGE gels were subsequently cut into two pieces, the upper (high MW part) was stained in Coomassie G250 to assess protein loading. The lower part was transferred to nitrocellulose membranes (midi, Bio-rad), using a Biorad Turboblot machine (Mixed MW, 7 min 25V). Membranes were appropriately cut and washed in 50 mL tubes using 5% BSA in TBST (0.1% Tween-20) for 1 hour at RT. Subsequently, membranes were incubated using primary antibodies against ABHD6 (Cell Signaling, D3C8N, 1:1000) in 5% BSA in TBST overnight at 4 °C. Next day, membrane were washed three times in TBST, and subsequently incubated with the appropriate secondary antibodies coupled to HRP (Santa Cruz). Development of the luminescence was performed using a chemiluminescence kit (BioRad), and scanned on a Chemidoc MP. The gel and blot images were further analyzed with Image lab software (v6.1, BioRad). Bands were quantified and the relative intensity was normalized on the Coomassie staining for protein loading normalization.

### RNA-sequencing data preparation

RNA sequencing data was obtained as described previously [30]. Raw counts were loaded into R using an EdgeR-voom-limma pipeline. The counts were loaded into a DGEList object and subsequently normalized using the trimmed-mean method (TMM) from EdgeR (v 4.2.1). Counts were equally distributed. Genes were filtered for high expression using the cpmbygroup function. Only genes with a cpm corresponding to a minimum of 10 counts, in at least one group (control, AD or resilient) were kept. This delivered 18,158 genes in 35 samples for further processing. Log2 CPMs were calculated using edgeR’s cpm() function with log set to TRUE. Subsequently, the log2 CPM were adjusted for sex and batch using the RemoveBatchEffect() function from Limma.

### Multi-omics factor analysis (MOFA)

Multi-omics factor analysis (MOFA) was performed using the R package MOFA2 (v1.8.0) using a python dependency with Basilisk 1.10.2. The input data was the full ABPP data (82 enzymes), the full lipidomics data (589 lipids) and a restricted RNAseq dataset. The RNAseq data was restricted based on variance, highly variable genes were selected by using a cutoff of 1.5 * mean log2 standard deviation. In addition, only protein-coding genes were kept. This yielded 1,699 genes for the input of MOFA. The MOFA model was trained with largely default settings. Scale_views was set to TRUE, to avoid excessively variant genes overruling more subtle changes in the lipidomics data. Convergence mode was set to “medium” and maximum iterations were 2000.

Derived factors were filtered based on minimum variance explained: we kept only those factors which explained at least 5% variance in at least one data modality. This led to 6 factors being kept for further analysis. 5 out of 6 Factors were normally distributed as tested by Shapiro-tests (p values > 0.05). Factors were then tested across groups using two-sided student’s t-test with BH multiple testing correction.

Biological pathways associated with the MOFA factors were determined with fgsea (1.30.0) and clusterprofiler (4.12.0), using the feature weights to rank the genes or lipids. Genes were related to gene ontology (“biological process”), lipids were grouped into lipid classes as described above. Factors were correlated (Pearson) to cell type abundance based on deconvoluted RNA-seq. data and to plaque and phosphorylated tau load from previously published data [30].

### Immunohistochemistry and immunofluorescence

Adjacent tissue used for the lipidomics and ABPP was used to cut 10 µm cryosections on a cryostat and was stored at –80 °C until use. For the Oil red O staining, 2 sections per donor were brought to room temperature and fixed in 4% PFA for 15 min. Sections were washed twice in 60% isopropanol for 5 min and then incubated in Oil red O (Abcam, lot number) for 30 min at 60 °C. Sections were rinsed in isopropanol, water and stained with haematoxylin for 30 s. The slides were coverslipped with Mowiol 4-88 mounting medium. Imaging of the sections was performed with an Axioscan Z1 (ZEISS, Oberkochen, Germany) at 20x magnification. Oil red O signal was quantified using Qupath (v0.5.0). Per section, two regions of interest (ROIs) were outlined covering all cortical layers of the grey matter. Within the ROI, a signal with twice the optical density of the background was considered Oil red O signal. The OD was multiplied by the surface area of the total Oil red O signal divided by the total area of the ROI. Within the same ROI, the cell counter function of Qupath was used on the haematoxylin signal after which positive cells were automatically counted based on the Oil red O signal.

For the immunofluorescent staining FFPE tissue from the SFG of the same donors was used. Briefly, FFPE sections were deparaffinized in xylene and rehydrated graded ethanol series. Antigen retrieval was performed using citrate buffer at pH 6 in a microwave at 800W for 10 min. The antibodies against PLIN2 (Progen Biotechnik; 1:200), Ionized calcium-binding adapter molecule 1 (Iba1, WAKO; 1:250), glial fibrillary acidic protein (GFAP, Cy3 conjugated, Sigma-Aldrich, 1:500) and NeuN (Millipore; 1:500) were diluted in diluent buffer (1%BSA, 0.5% triton in TBS) overnight at 4 °C. Secondary antibodies (donkey anti-mouse Cy3, donkey anti-guinea pig 488, donkey anti-rabbit Cy5, ThermoFisher Scientific; 1:400) were diluted in the diluent buffer and incubated at room temperature for 1 hour. Sections were treated with 0.1% sudan black B in 70% EtOH for 5 min and nuclei were visualized with DAPI. Sections were coverslipped with Mowiol 4-88. Representative pictures were taken with a Leica TSC SP5 microscope (60x magnification, 1056×1056 and 100 Hz).

## Results

To study alterations in lipid metabolism, we first analysed the lipid content of the grey matter in the superior frontal gyrus (SFG) from 12 control, AD and resilient individuals (Table 1), using targeted LC/MS-MS. We analysed 629 lipid species across 31 lipid classes, covering a wide range of lipid classes including phospholipids, plasmalogens, sphingomyelins, ceramides, bismonoacylglycerolphosphates, acyl-glycerides, free fatty acids and their oxidized derivatives (oxylipins). An unbiased analysis of the main sources of variance using principal component analysis (PCA) showed that most variance in the lipidome did not relate to disease classification (Figure 1a-b). The first two components also did not relate to donor characteristics like age, post-mortem delay, pH of the CSF or sex, indicating large sources of variability between individuals for currently unknown reasons (Figure 1a). Relating the first 10 principal components to disease characteristics did, however, show that PC3 and PC7 correlated with AD pathology. A plot of these components separated controls from AD, with resilient individuals overlapping with both groups (Figure 1c).

**Figure 1.**
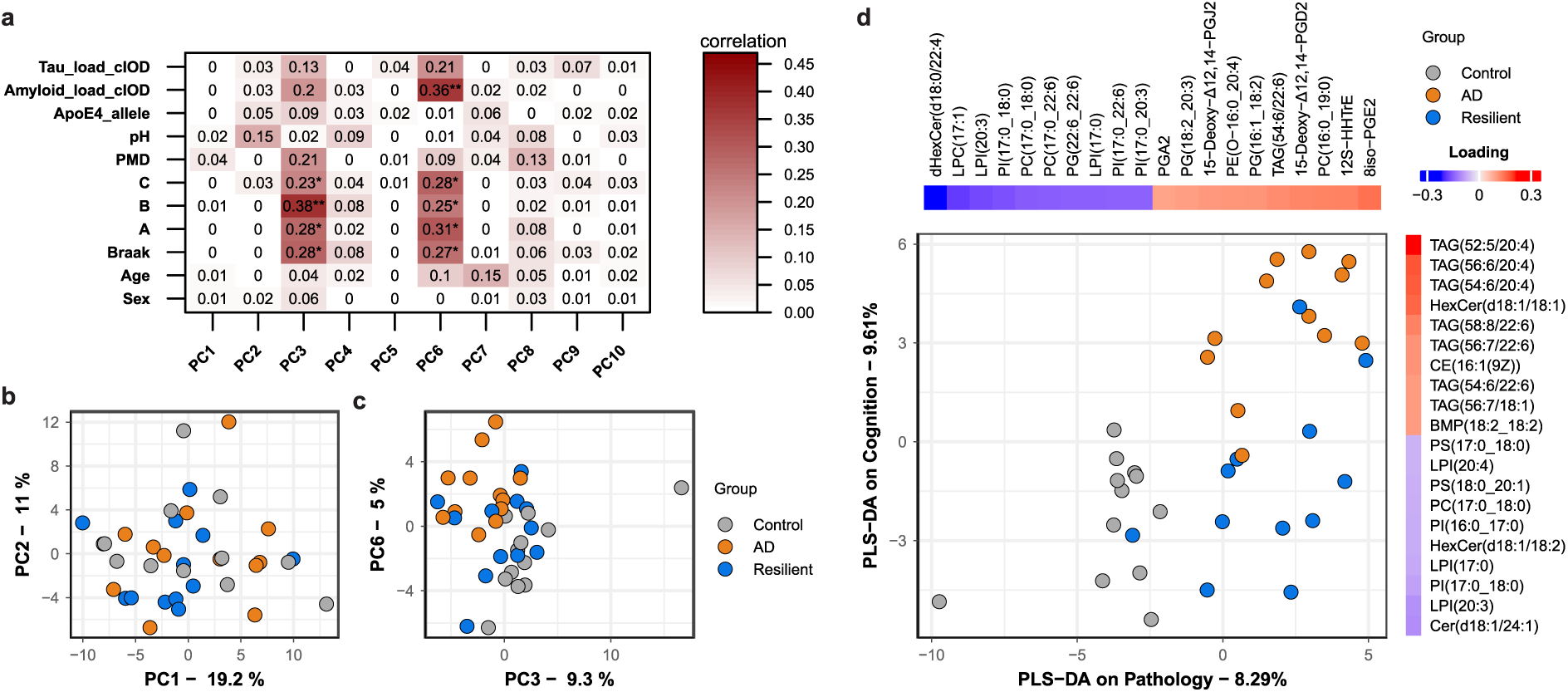
Heterogeneous lipidomic data can be separated by group based on TAGs and oxylipins. a. Correlations of metadata and the first 10 principal components (PCs). b. PCA plot of PC1 and PC2 does not separate groups. c. PCA plot of PC3 and PC7 shows separation between groups. d. Partial least squares-discriminant analysis (PLS-DA) shows separation of groups mainly driven by TAGs and oxylipins. The loadings of the top 10 positive and negative loadings are plotted alongside their respective axis.

Partial least-squares discriminant analysis (PLS-DA) was used to identify components that best separated the groups. As donors were selected based on their amount of pathology and cognition, we used those parameters as input for PLS-DA. The neuropathological scores were significantly different between AD and resilient individuals versus control donors, while there was no difference between AD versus resilient individuals (Table 1). PLS-DA on AD pathology and cognition revealed that oxylipins were driving the separation between controls and AD and resilient individuals, while triacylglycerols (TAGs) contributed to the difference between controls and resilient donors from demented individuals, albeit only explaining a small percentage of the variance.

### Inflammatory oxylipins from ω6 free fatty acids are upregulated in response to pathology

To directly identify lipids that were significantly different between groups, we analysed differentially abundant lipids applying a Benjamini-Hochberg (BH) correction with a 25% false discovery rate (FDR) due to high non-disease-related variance. The fold changes of both AD and resilient groups compared to controls correlated strongly with each other (Figure 2d, Pearson’s correlation coefficient R = 0.65, p<2.2E-16), indicating a similar lipidomic response to AD pathology. Oxylipins, products of free fatty acid oxidation and markers of neuroinflammation, showed the most changes between AD patients and controls (Figure 2a). In resilient individuals, oxylipins trended to increase, but this did not reach statistical significance (Figure 2b,d).

**Figure 2.**
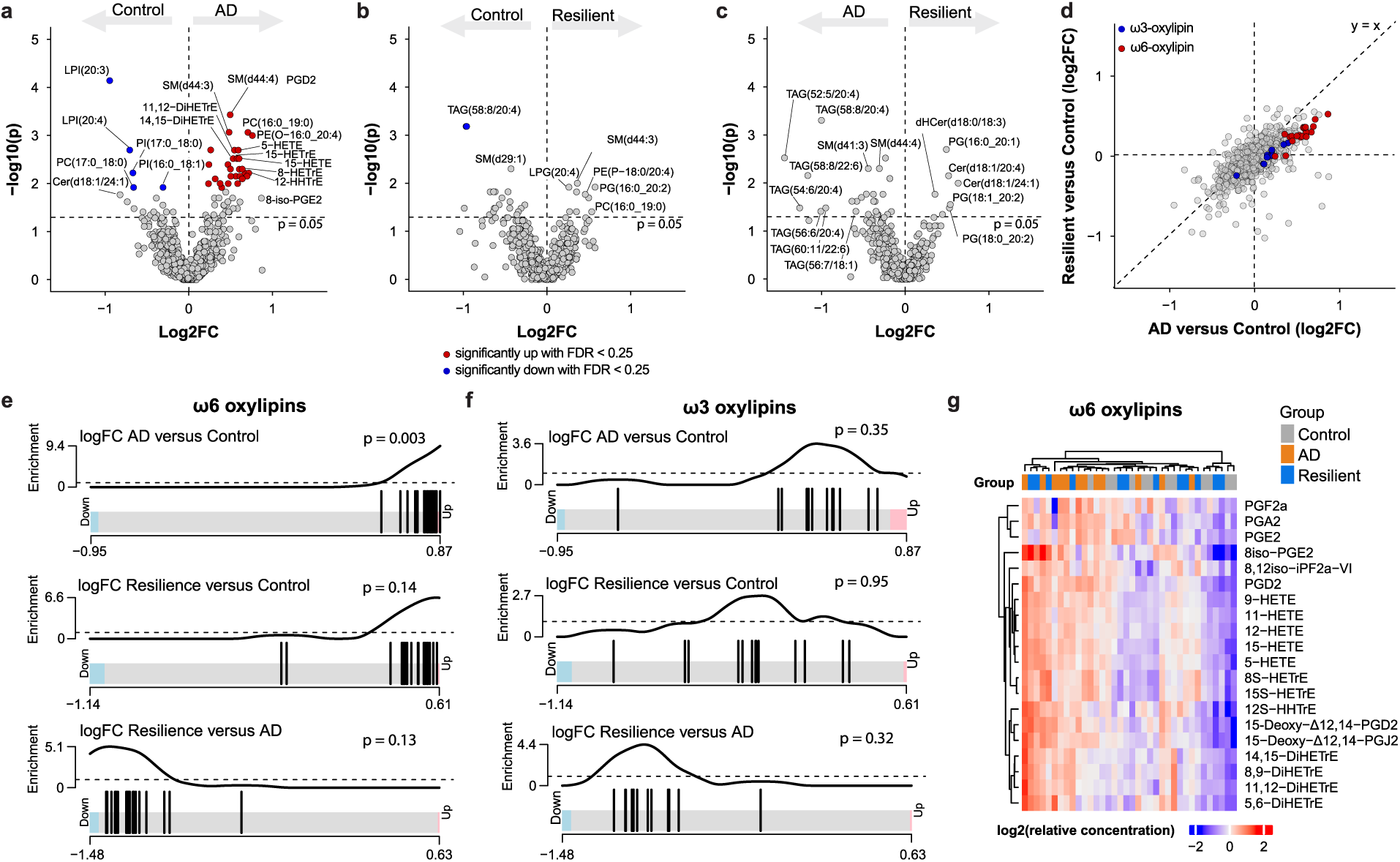
ω6-oxylipins are increased in AD donors. a-c. Differentially abundant lipids between the three groups in this study. p-values were calculated by Wilcoxon rank sum tests. The unadjusted p-value is plotted. Benjamini-Hochberg multiple testing correction was applied to identify those lipids that are significant with a false discovery rate of 25% (red, significantly up; blue, significantly down). d. Quadrant plot of fold-changes from AD versus control and resilient versus control. ω3-and ω6-oxylipins are indicated in blue and red, respectively. e-f. barcode plots of enrichment analysis using ROAST shows that ω6 oxylipins (e) but not ω3 oxylipins (f) are enriched in AD. g. Heatmap of normalized ω6-oxylipin levels, presented as z-scores, shows that these are generally higher in the AD donors.

Notably, only ω6-derived oxylipins, not ω3, showed positive fold changes (Figure 2d). The enrichment of ω6-derived oxylipins was confirmed by rotation gene-set testing (ROAST) in the AD versus control groups (p = 0.003), but not in the resilient individuals versus controls (p = 0.14). ω3-oxylipins were not enriched in any comparison, suggesting that the increase in oxylipins in AD is selective for ω6 free fatty acids (Figure 2f). This may be due to decreased ω3-oxylipin precursors docosahexaenoic acid (DHA, 22:6-ω3) and eicosapentaenoic acid (EPA, 20:5-ω3) (Figure S1a,b). No changes were found in arachidonic acid (AA, 20:4-ω6) and dihomo-gamma-linolenic acid (DGLA, 20:3-ω6) levels, precursors for the ω6-oxylipins (Figure S1c,d). Taken together, these data show that ω6-oxylipins are more abundant in the cortex of AD donors and follow a similar trend in resilient donors.

### Lower TAGs levels and lipid droplets in resilient individuals

Because we observed a trend for several species of triacylglycerols (TAGs) to be reduced in resilience compared to AD (Figure 2c), and TAGs associated with cognition in the PLS-DA (Figure 1d), we further investigated TAG levels across the groups. We observed a separation of AD and resilient donors by hierarchical clustering (Figure 3a). A subset of TAGs showed negative fold changes in the resilient group, while they had positive fold changes in the AD group (Figure 3b). These changes were more pronounced in TAGs with a higher degree of unsaturation, while more saturated TAGs remained generally unchanged (Figure 3c). Enrichment analysis revealed a significant decrease in TAG levels in resilient donors compared to AD (p = 0.045, Figure 3d), suggesting lower total TAG levels in the resilient donors. Since TAGs are major components of lipid droplets (LDs), these data may reflect alterations in lipid droplets in resilient individuals compared to AD patients.

**Figure 3.**
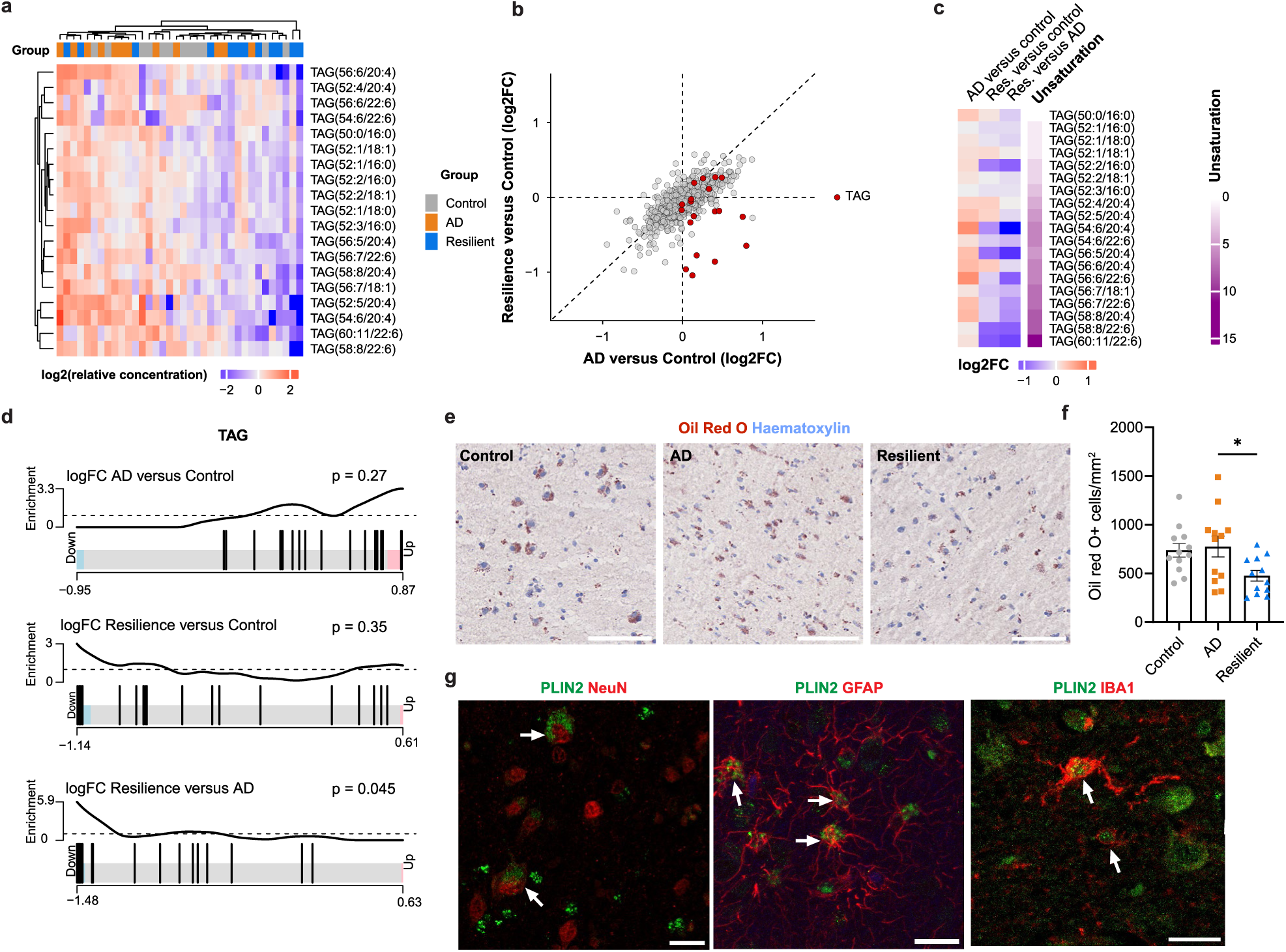
TAGs and lipid droplets are reduced in resilient donors. a. Heatmap of normalized TAG levels shows separation of the resilient from AD donors. b. Quadrant plot of foldchanges from AD versus control and resilient versus control shows some TAGs are less abundant in the resilient group, while more abundant in the AD group. c. Heatmap of fold changes of the different TAGs shows that mainly the more unsaturated TAGs are less abundant in the resilient group. d. Enrichment analysis using ROAST shows that TAGs are less abundant in the resilient group. e. Representative Oil red O (ORO) of a resilient donor staining in the superior frontal gyrus of the different groups. f. Quantification of ORO positive cells in the cortex across groups. Data are represented as mean ± SEM. * p < 0.05, ANOVA with Tukey post-hoc. g. PLIN2, a marker for lipid droplets, colocalizes with GFAP-, NeuN-and Iba1-positive cells. Representative pictures are from the same AD donor. Scale bars in panel e are 100 µm and in g 20 µm.

To obtain an independent line of evidence for alterations in lipid droplet content between resilient and AD donors, neutral lipids were visualized in adjacent tissue sections using Oil Red O (ORO) staining. A higher number of cytoplasmic LDs was observed in cortical cells of AD donors compared to resilient donors (Figure 3e-f: F=4.09, p=0.027; AD vs control, p=0.942; resilient vs control, p=0.071; AD vs resilient, p=0.034). No significant change was observed between AD and control donors. We did not find an association of both TAGs and lipid droplets with ApoE4 genotype (Figure S2). Staining of PLIN2, an integral lipid droplet membrane protein marker, revealed that astrocytes, microglia and neurons across all donor groups contained lipid droplets (Figure 3g), suggesting that the changes in LDs are not limited to one specific cell type.

### Activity-based protein profiling uncovers changes in enzyme activities in response to pathology

To investigate which lipid metabolizing enzymes were potentially involved in the changes in oxylipins and TAGs, we performed activity-based protein profiling (ABPP), a technique to enrich active enzymes from native tissues using chemical probes [40]. Specifically, we used fluorophosphonate and β-lactone probes to target serine and cysteine hydrolases, which are intricately involved in lipid metabolism in the brain [41].

We quantified relative enzyme activities for 76 enzymes. The activity of these did not correlate with the expression of their respective genes, possibly due to post-transcriptional and post-translational regulation (Figure S3). Consistent with the lipidomics dataset, we observed small changes in enzymatic activities across groups. Tripeptidyl peptidase 2 (TPP2) was the only enzyme differentially altered between control and AD donors (Figure 4a), while alpha/beta-Hydrolase domain containing 6 (ABHD6), ubiquitin C-Terminal Hydrolase L1 (UCHL1), 4-aminobutyrate aminotransferase (ABAT) or platelet activating factor acetyl hydrolase 1b catalytic subunit 3 (PAFAH1B3) were altered between control and resilient donors (Figure 4b). No changes were observed between AD and resilient donors (Figure 4c). Most fold changes were similar between resilience and AD groups (Figure 4d), indicating that enzyme activities mainly associate to AD pathology.

**Figure 4.**
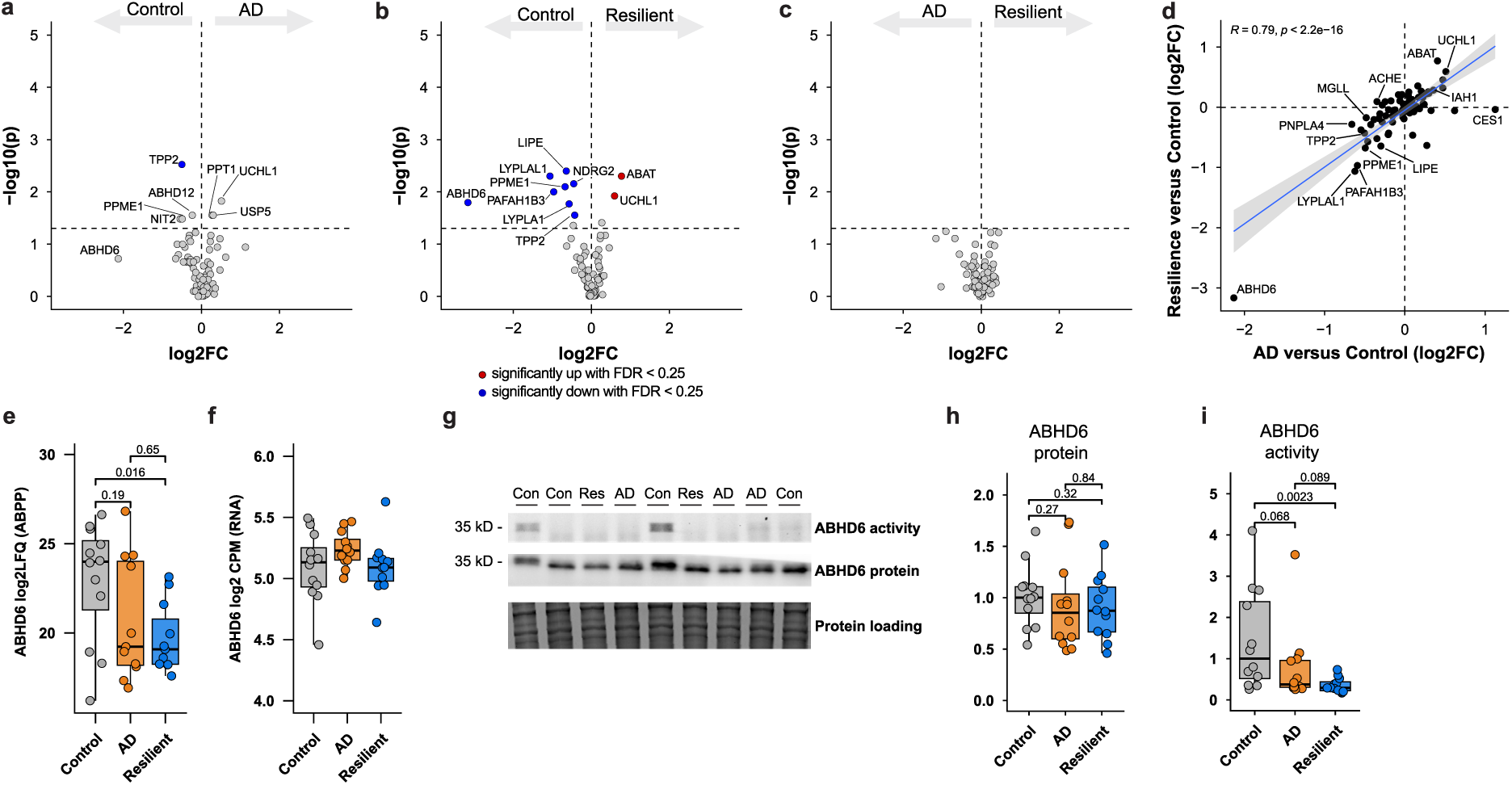
Activity-based protein profiling shows similar changes in enzyme activities between AD and resilient donors. a-c. Differentially active enzymes between the three groups in this study. p-values were calculated by Wilcoxon rank sum tests. Benjamini-Hochberg multiple testing correction was applied to identify those enzymes that are significant with a false discovery rate of 25%. d. Quadrant plot of fold changes from ABPP results between AD versus control and resilient versus control show that the changes in resilient and AD donors are highly correlated. e. ABHD6 levels from the ABPP between the groups, showing lower activity levels of ABHD6 in the resilient group. p-values are the raw p-value from a-c. f. There are no differences in mRNA levels of *ABHD6*. Transcript levels are derived from a previously generated bulk RNA-seq. dataset [30]. Representative gels of ABHD6 activity by fluorescent probe LEI-612-BDP-TMR, and ABHD6 protein expression by western blot across the three groups. Full gels can be found in Figure S3. h. Quantified protein levels of ABHD6 from panel g by western blot. i. Quantified activity levels from panel g. p-values represent Wilcoxon rank sum tests (N=12 for each group).

Notably, ABHD6 was the most downregulated enzyme in the cortex of resilient patients compared to control donors, with a similar trend in AD donors (Figure 4d,e). Remarkably, ABHD6 activity varied with several orders of magnitude across donors for unknown reasons, while *ABHD6* mRNA expression showed no differences between groups and a much smaller variability [30] (Figure 4f), suggesting a disconnection between protein activity and gene expression. Western blot analysis confirmed that ABHD6 protein levels were consistent across all samples, while gel-based ABPP analysis revealed that ABHD6 activity was significantly lower in resilient individuals, with a negative trend in AD donors (Figure 4j: resilient vs control, p=0.0023; AD vs control, p=0.068, AD vs resilient, p=0.089). Overall, these results show that decreased ABHD6 activity is associated to AD pathology in a manner that is independent of its expression in both resilient and AD donors.

### Multi-omics data integration couples oxylipin levels to loss of inhibitory neurons in AD and resilience

Finally, to link the lipid and enzyme activity patterns with the global transcriptome and cell types, we integrated three datasets using multi-omics factor analysis (MOFA) [42]. We trained the MOFA model on the lipidomics (N= 36, 589 lipids) and ABPP (N=36, 82 enzymes) data, together with the most variable genes from a previously generated transcriptome dataset (N=35, 1,699 genes) [30] (Figure 5a, S5). We limited the derived factors to those explaining at least 5% variability in one data modality. Six orthogonal factors were identified, each capturing independent variance across the datasets (Figure 5b, S5).

**Figure 5.**
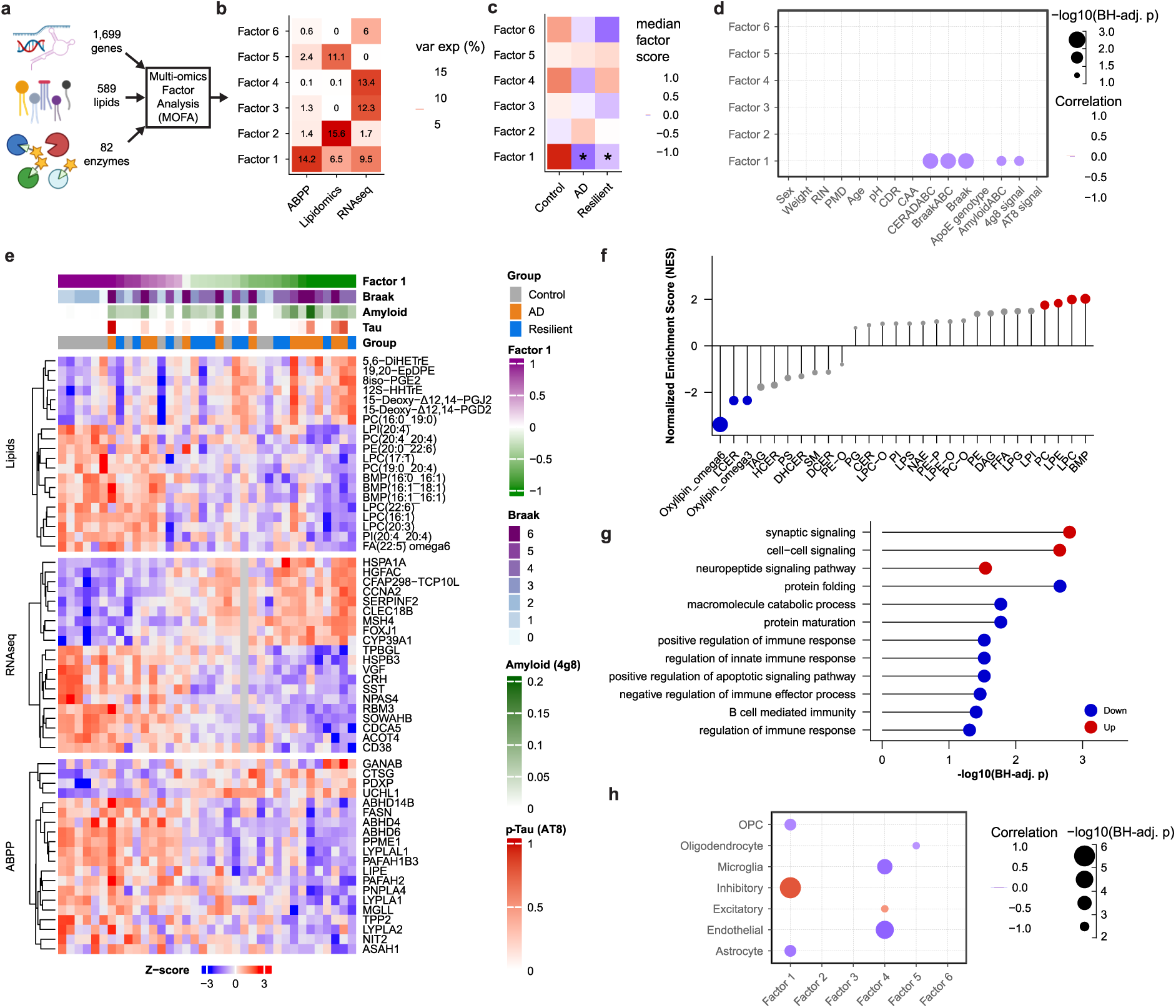
A multi-omics response in donors with AD pathology that is associated with Aβ plaque levels. a. Input data of the different data modalities for the MOFA analysis. b. MOFA identified 6 factors that each explained at least 5% of the variance in at least one data modality. c. median factor scores per group, *q < 0.05, student’s t-test with BH-correction. d. Correlation (Pearson) plot of donor characteristics and pathological parameters with the MOFA factors. e. Heatmap of the top-20 variables associated with Factor 1 from all three data modalities. Data is depicted as z-scores. Legends indicate per sample the group, quantified p-tau pathology (AT8) and Aβ plaques (4g8), Braak stage and Factor 1 score. f. Enrichment analysis of lipid species associated with factor 1 shows that mainly ω6-oxylipins associate with factor 1. Data is plotted as normalized enrichment scores (NES). g. Gene-set enrichment analysis of genes associated with Factor 1. Data is represented as the BH-adjusted p-value for enrichment of GO terms positively (red) or negatively (blue) associated with Factor 1. Only Go-terms with q<0.05 are shown. h. Correlation of MOFA factors with cell type abundances determined by deconvolution of RNAseq data. Data represent Pearsons correlations with BH-correction for multiple testing.

Only factor 1 displayed significant association with the three groups, being significantly lower in both AD and resilient donors (Figure 5c). It captured variance from all three data sources (Figure 5b). Factor 1 was significantly correlated with neuropathological scores for AD (Figure 5d), specifically showing a negative correlation with quantified Aβ plaques (by 4g8), but not with phosphorylated tau (AT8). This suggests that factor 1 reflects a response to Aβ plaques. Factor 1 did not relate to age, post-mortem delay or other disease unrelated parameters, showing that it is a neuropathology specific multi-omics response (Figure 5d).

The genes, lipids and enzymes with the highest loadings of factor 1 included oxylipins, heat shock protein-related genes, *SST* and *VGF*, and MGLL and ABHD6 (Figure 4f, S6). Enrichment analysis using the lipid and gene loadings indicated that ω6-oxylipins were negatively associated with factor 1, capturing their upregulation in response to Aβ pathology (Figure 5f). In addition, genes related to synaptic signalling were positively associated with factor 1, while genes related to protein folding and immune responses were negatively correlated (Figure 5g, S7). This suggests that factor 1 reflects a loss of synaptic signalling and increased immune responses and heat shock proteins in the AD and resilient cortex, in relation to Aβ plaque load.

Next, we correlated the MOFA factors to cell type abundances based on cell type deconvolution of the transcriptomics data [30]. The proportion of astrocytes and oligodendrocyte precursor cells (OPCs) were negatively correlated to factor 1, possibly indicating gliosis. In contrast, inhibitory neurons showed a strong positive correlation with Factor 1 (R = 0.78, *p* = 4.6E-8, Figure 5h) indicating that increased inflammatory ω6-oxylipins associate with loss of inhibitory neurons in both AD-and resilient donors. In conclusion, multi-omics data integration captured a response to Aβ plaques, characterized by inflammatory ω6-oxylipins, gliosis, a loss of inhibitory neurons and reduced synaptic signalling.

## Discussion

How alterations in lipid metabolism and signalling might contribute to the pathophysiology of AD is so far not well understood. By using a multi-omics approach, we provide evidence for alterations in inflammatory ω6-oxylipins, loss of inhibitory neurons and a decrease in synaptic signalling that are associated with Aβ plaques. Overall, whereas most changes in lipid metabolism were similar in the AD and resilient groups, we found lower levels of TAGs and lipid droplets in the resilient donors specifically.

Increased TAGs and cholesterol esters, major components of lipid droplets, were previously found close to plaques in AD, particularly in glial cells [16, 43]. Increased lipid droplets in glial cells have been associated with more proinflammatory states [16, 17] and in animal models microglia form lipid droplets after Aβ exposure, which is accompanied by a decrease in Aβ phagocytosis [44]. Green *et al.* recently reported lipid-laden microglial states in AD, associated with the development of AD pathology [8]. Our data could indicate that such microglial states might be reduced in resilient donors. As these lipid-laden microglial states might be more proinflammatory [45], a reduction of proinflammatory microglial subtypes may be expected in resilient donors, which is in line with previous research [10, 46, 47]. In addition, a reduced lipid load in microglia was recently described to enhance amyloid plaque clearance in a mouse model of AD [49]. While others have shown that lipid droplets are strongly associated to the ApoE4 genotype [16, 43, 51], in our data TAGs and lipid droplets did not correlate with ApoE4. In the current study, TAGs and lipid droplets were measured across all cell types, which may explain why in our results there is no correlation with ApoE4 genotype. In addition, donors included in our study predominantly had heterozygous ApoE4, whereas previous studies primarily related to biallelic ApoE4. Taken together, the alterations in lipid droplets between the resilient and AD donors may reflect a different glial response to pathology in resilience compared to AD.

The lipid class with the most significantly altered lipids was the ω6-oxylipins, which were also found in the multi-omics response and are in line with previous observations [48]. Oxylipins have been associated with inflammatory processes in AD, in which ω6-derived oxylipins are generally considered pro-inflammatory, while ω3-derived oxylipins can reduce neuroinflammation [50, 52]. Another class of closely associated lipids, the specialized pro-resolving mediators (SPMs) also have important roles in protecting against neuroinflammation. Although they were part of our targeted lipidomics platform, we did not detect these lipids, most likely due to their low concentration. As mainly ω6-derived oxylipins were enriched in AD donors and in MOFA factor 1, this may indicate increased levels of neuroinflammation in these donors. Interestingly, mainly unsaturated TAGs were more abundant in AD patients compared to resilient donors, which can serve as precursors for oxylipins. This could suggest that the reduced TAGs in the resilient donors possibly have resulted in lower levels of oxylipins. The ω6-oxylipin response in the cortex of AD donors indeed was stronger than in resilient donors. Furthermore, MOFA factor 1 correlated with Aβ plaques but not with phosphorylated tau, indicating that processes captured by this factor are possibly a specific response to Aβ plaques. This response may be similar in both AD and resilient donors, suggesting that this could either be a general response or a response associated with early stages of AD pathophysiology, rather than related to diminished cognition, as the resilient donors are cognitively intact. Early features of AD are synaptic loss [53, 54] and loss of inhibitory interneurons [55–58], possibly driven by Aβ-induced neuroinflammation. Our data indicate that these processes might already be ongoing in the cortex of resilient individuals, despite having intact cognition.

The enzyme activities determined by ABPP did not reveal enzymes that could explain the difference in TAGs and lipid droplets in this cohort. Most enzyme activities changed similarly in both the resilient and AD donors compared to controls, indicating that this group of enzymes might be altered as a general reaction to neuropathology. We further profiled the mono-acyl glycerol hydrolysing enzyme ABHD6. ABHD6 was significantly reduced in resilient donors and showed a trend towards lower levels in AD. ABHD6 is involved in the hydrolysis of 2-AG to AA [59, 60] and is reduced in hippocampal neurons in AD patients [26]. Besides its role in lipid metabolism, ABHD6 is also involved in α-amino-3-hydroxy-5-methyl-4-isoxazolepropionic acid receptor (AMPAR) trafficking [61]. Strikingly, ABHD6 activity was uncoupled from its expression levels, which, to our knowledge, has not been reported so far. One possibility might be that specific point mutations in these donors affect ABHD6 activity but not its protein levels. Alternatively, post-translational regulation could cause a disconnection between protein levels and its activity. In addition, UCHL1 activity levels were higher in resilient donors and showed a trend in the same direction for the AD donors, which has been linked to lower levels of APP and amyloid-β [62]. TPP2 activity was significantly reduced in both resilient donors and AD patients. TPP2 is involved in antigen processing, cholecystokinin degradation, cell growth, and DNA damage repair [63–65], and lower levels have been linked to cognitive deficits in mice [66]. Both enzymes showed a similar direction in the AD and resilient donors, indicating that this could be a general reaction to AD pathology. Interestingly, others have shown that higher levels of the protein PAFAH1B3 might be associated with resilience [67, 68]. In the ABPP data, PAFAH1B3 was lower in the resilient donors compared to the control donors. It plays an important role in brain development [69], and its paralog PAFAH1B2 has been shown to reduce Aβ peptide production [70]. The difference with the current study may be explained by the difference in measuring protein abundance with proteomics or activity with ABPP.

There was significant disease unrelated variability in both the lipidomic and ABPP datasets, highlighting the need for many donors to be included in omics-studies like the current study. However, the donors used here have previously been comprehensively characterized to minimize confounding factors such as latent or comorbid disease [30]. This approach ensured the inclusion of resilient donors and excluded a large number of preclinical AD donors, which are often included by others [67], and of which it is uncertain if they would become resilient and postpone the onset of AD or develop dementia sooner if pathology would progress. Despite having 12 donors for each group, our study appeared underpowered to quantify the subtle changes in lipid composition with more statistical certainty. Furthermore, it might be possible that the bulk approaches used in the current study are not sensitive enough to detect local differences. Many of the previously indicated roles of lipids in AD are microglia related, which are less abundant than other cell types. Furthermore, many changes could happen in close vicinity to plaques. Recently, spatial mass spectrometry approaches have shown increased phospholipid species around plaques in AD patients [71] and differences in gangliosides in Aβ plaques from AD patients compared to Aβ plaques found in controls [72]. Future studies should focus on whether changes in lipid metabolism in relation to AD are found in specific microglia populations or surrounding specific Aβ plaque types.

In conclusion, we show that there is a general response to Aβ plaques that is associated with an increase in ω6-oxylipins and loss of interneurons. Furthermore, we provide the first evidence that subtle differences occur in lipid metabolism between AD and resilient individuals, demonstrating lower levels of TAGs and lipid droplets in the resilient donors.

## Acknowledgements

We are grateful to the brain donors and their families for their commitment to the Netherlands Brain Bank donor program. Bogdan I. Florea is acknowledged for managing proteomics facilities and equipment.

## Author contributions

LV and DS performed the donor selection and additional diagnostics. DV, LV, XD, MM collected data. Data collection was supervised by TH, AH and MS. DV, LV, BJ, MP and MM analyzed data. LV and DV drafted and revised the manuscript and prepared the figures with input from all authors. LV, DV, IH, JV and MS designed and coordinated the study. All authors read and approved the final manuscript.

## Availability of data and materials

No custom code was developed for this study. Mass spectrometry data for activity-based protein profiling will be available through the ProteomeXchange Consortium via PRIDE upon publication with accession code PXD057993. RNA sequencing data used this study was obtained and previously reported by de Vries *et al.* [30] and is available through the Gene Expression Omnibus with accession number GSE261817. Other data and R code is available upon request to the corresponding authors.

## Ethics approval and consent to participate

Informed consent for a brain autopsy and for the use of the brain material and clinical data for research purposes was obtained by the NBB according to international ethical guidelines and were approved by the Medical Ethic Committee of the VU Medical Center, Amsterdam, the Netherlands.

## Consent for publication

Not applicable.

