## Supplementary Figures for "Reduced triacylglycerols and lipid droplets are associated with resilience to Alzheimer’s disease"

* these authors contributed equally

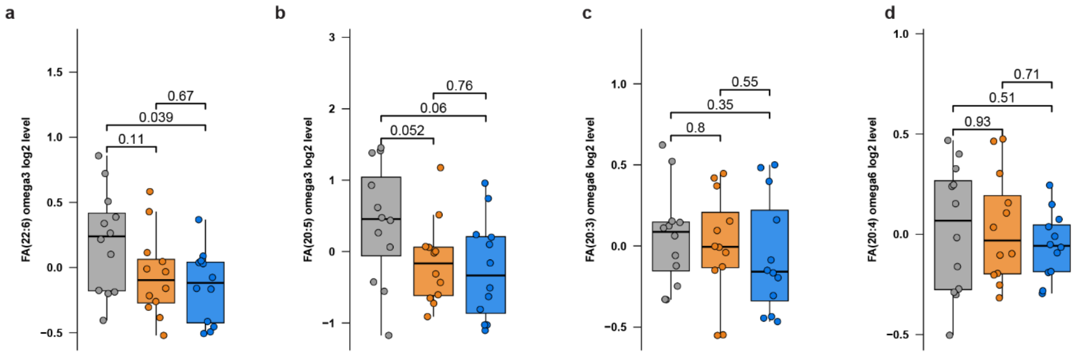


**Supplementary figure 1. a-d,** Relative levels of oxylipin precursors DHA (a, FA(22:6) omega 3), EPA (b, 20:5 omega 3), DGLA (c, FA(20:3) omega 6) and AA (d, FA(20:4) omega 6). Statistical test is Wilcoxon rank sum test without any adjustment for multiple testing.

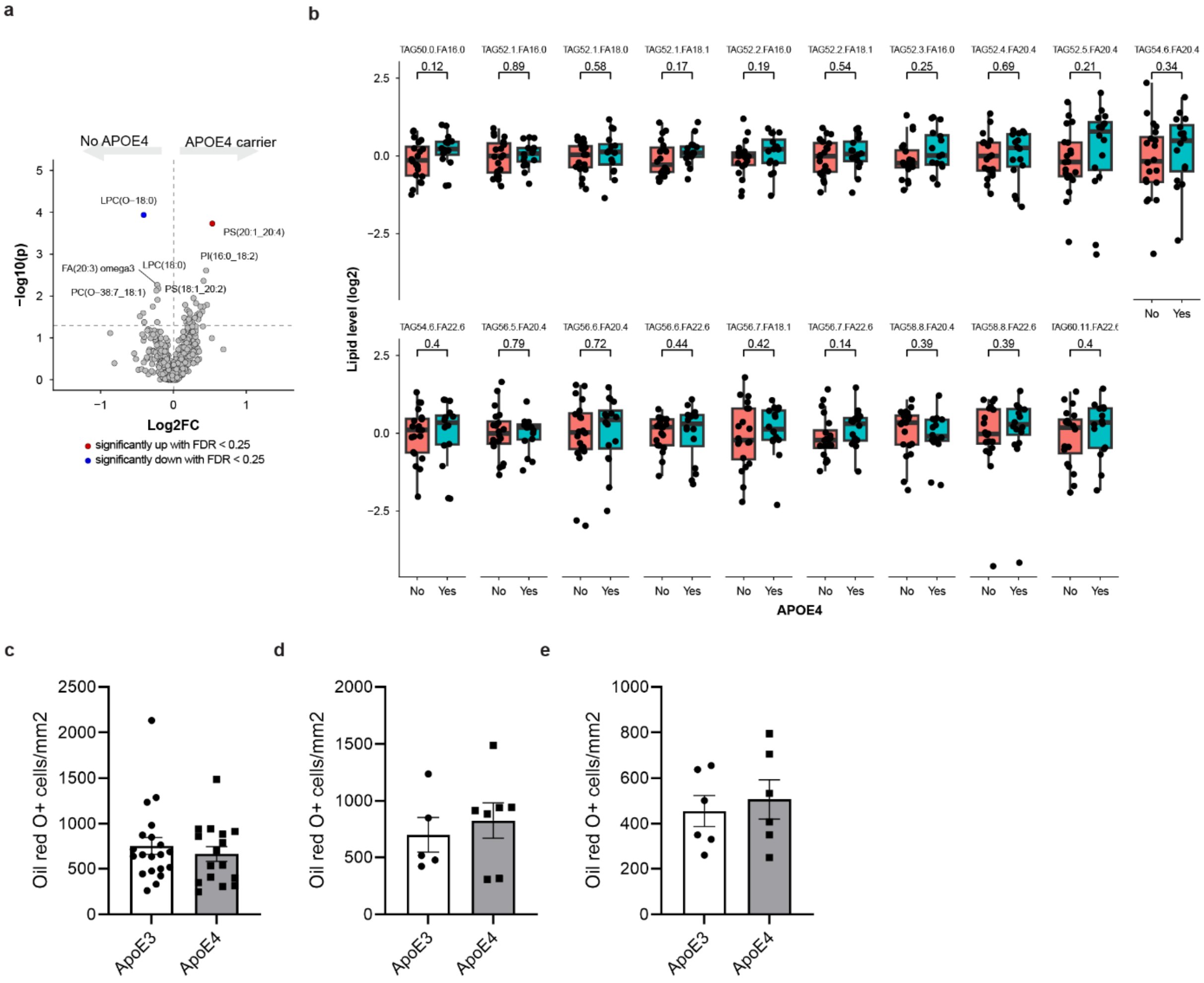


**Supplementary figure 2. APOE4 allele presence does not influence lipid droplet accumulation in this cohort.** APOE4 allele positive donors include 4/4 (N=1), 4/3 (N=14) and 4/2 (N=1) and APOE4 negative donors include 3/3 (N=17) and 3/2 (N=3) **a,** Volcanoplot of differentially expressed lipids between APOE4 postive and – negative donors. p-values were calculated by Wilcoxon rank sum tests. The raw p value is plotted. An FDR of 25% was applied to identify significantly altered lipids, indicated by color. **b,** boxplots of TAG levels between APOE4 positive and negative donors. p values represent Wilcoxon rank sum tests. **c-e,** Oil Red O positive (lipid droplet) positive cells by APOE4 genotype. No difference was observed in the whole cohort (c), the AD group (d) or the resilient group (e).

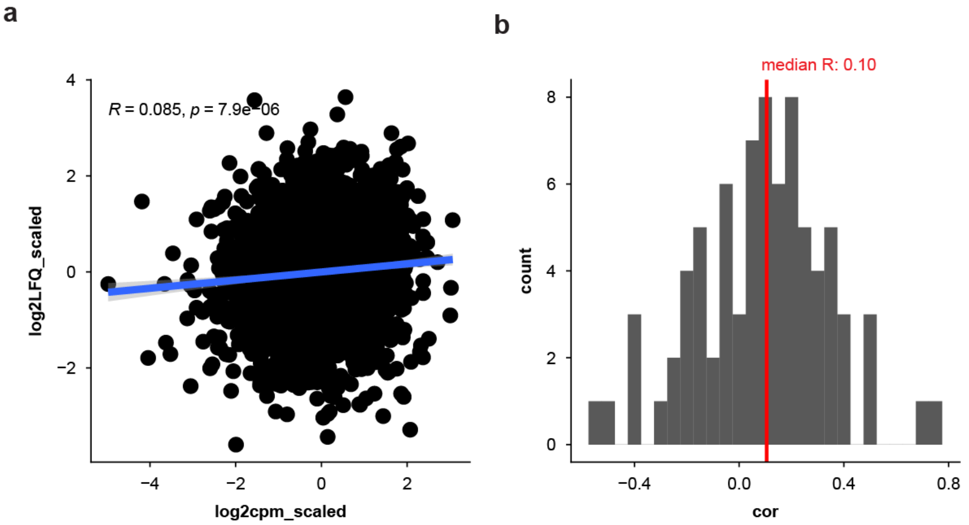


**Supplementary figure 3.** Correlation of enzyme activities with the expression of their genes. A. pooled analysis of all ABPP enzymes and their corresponding mRNA. Data was z-score and correlated using a Pearson correlation. b. histogram of the correlation coefficients of individual RNA-enzyme pairs. Data is plotted as Pearson's R and the median R is indicated.

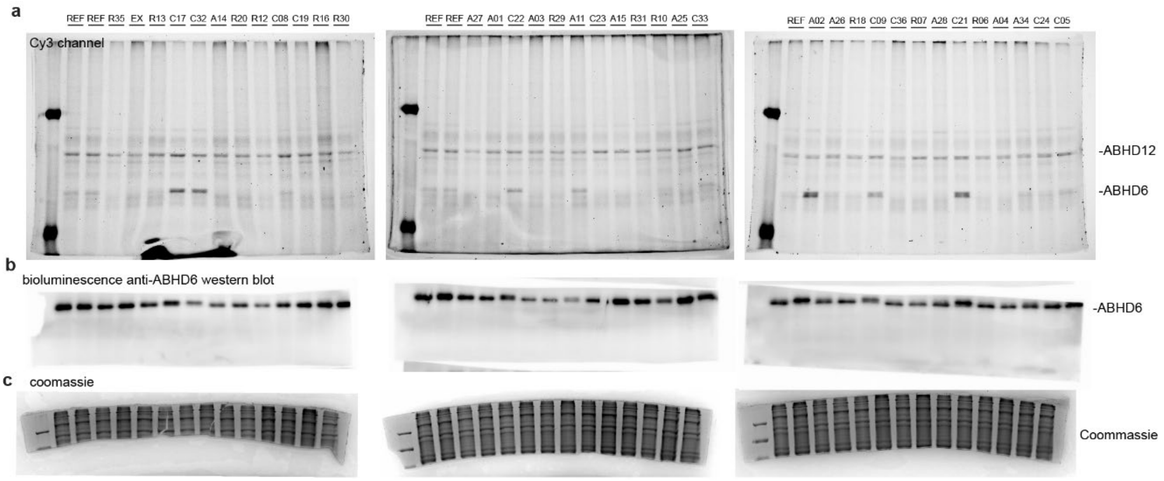


**Supplementary figure 4.** Full pictures of gel-based ABPP and ABHD6 Western blot. a. Bands of ABHD12 and ABHD6 activity using the probe LEI-612-BDP-TMR (200 nM). ABHD12 appears at ~55 kDa while ABHD6 appears as the variable bands at ~35kDa. b. Protein bands of ABHD6 determined by western blot. c. Full pictures of the Coomassie staining. Samples are labelled 1-36 with A=Alzheimer, R=resilient and C=control, ref=reference sample for quality control (not quantified), ex=excluded sample.

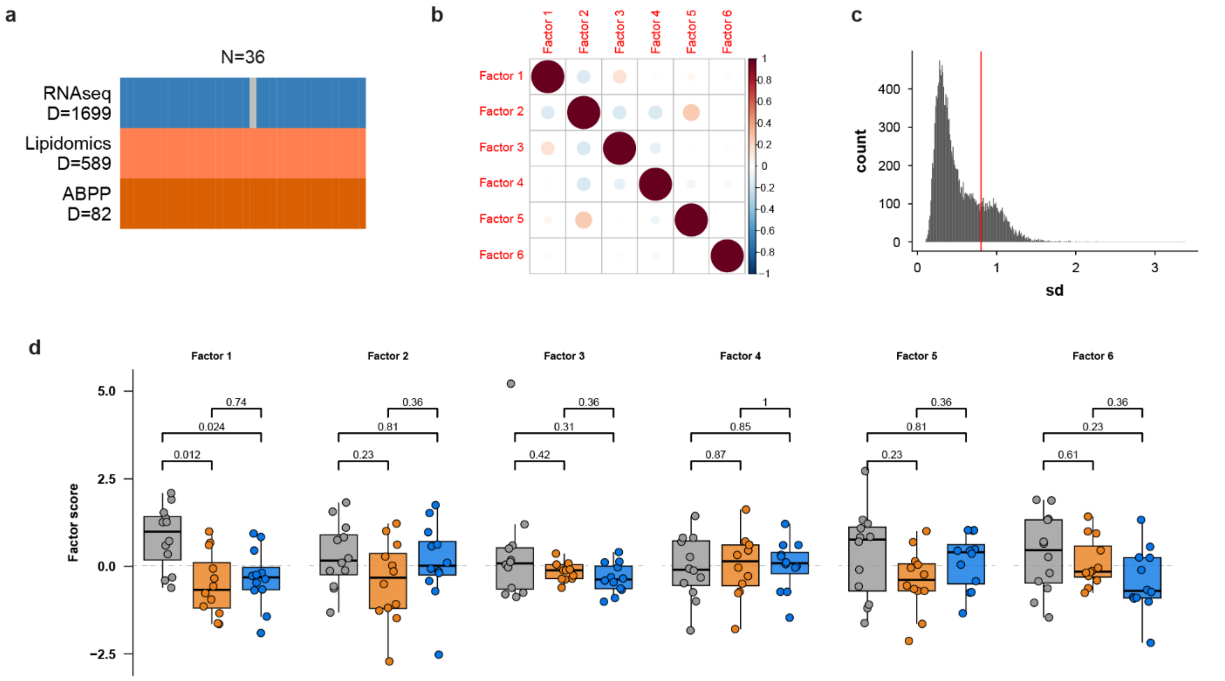


**Supplementary figure 5.** Six Factors derived by multi-omics data integration.

a. Input data for MOFA. b. MOFA factors are orthogonal, meaning that each factor captures unique variance from the datasets. c, transcriptomics data was restricted to the 1,699 most variable genes. The red line indicates the cutoff. d. All factors derived from the MOFA analysis tested across groups. p-values represent two-sided student's t-test with BH multiple testing correction (q-values).

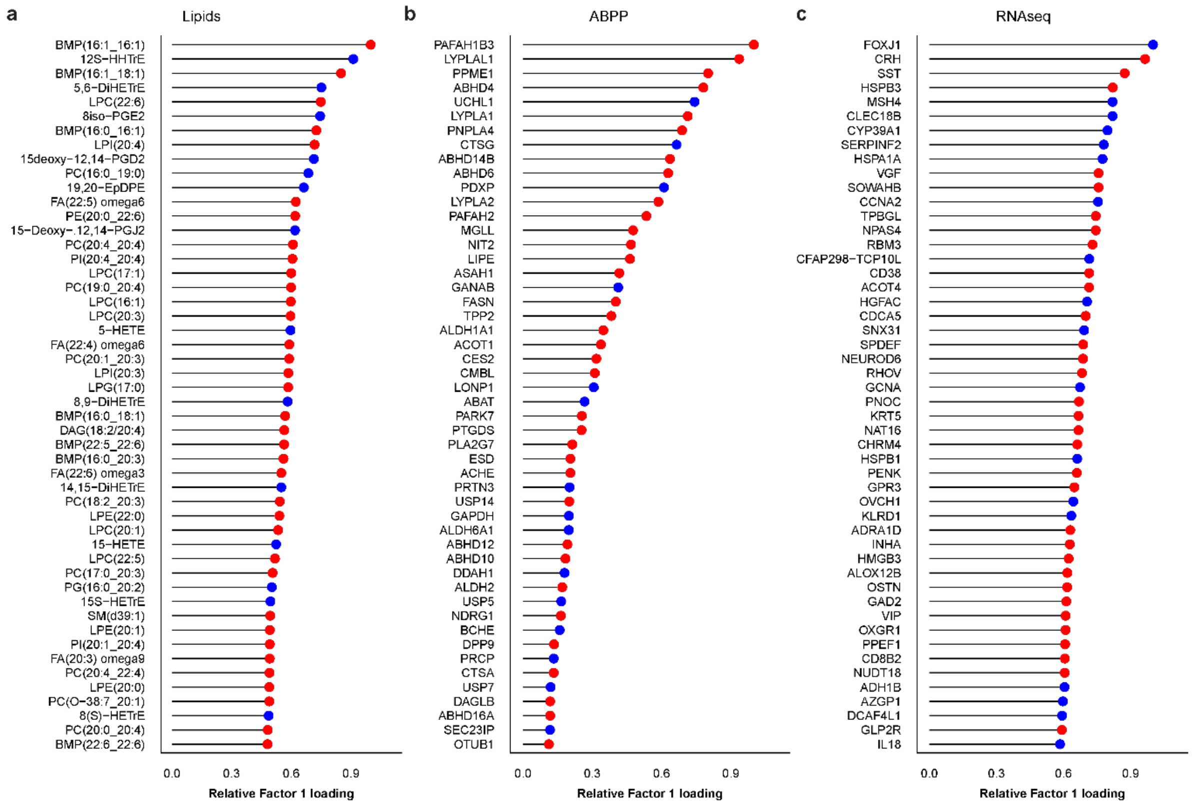


**Supplementary figure 6.** Top 50 weights of MOFA Factor 1 for lipids (a), enzymes (b) or genes (c). Colours indicate direction, where red means positive association with Factor 1, and blue means negative association to Factor 1.

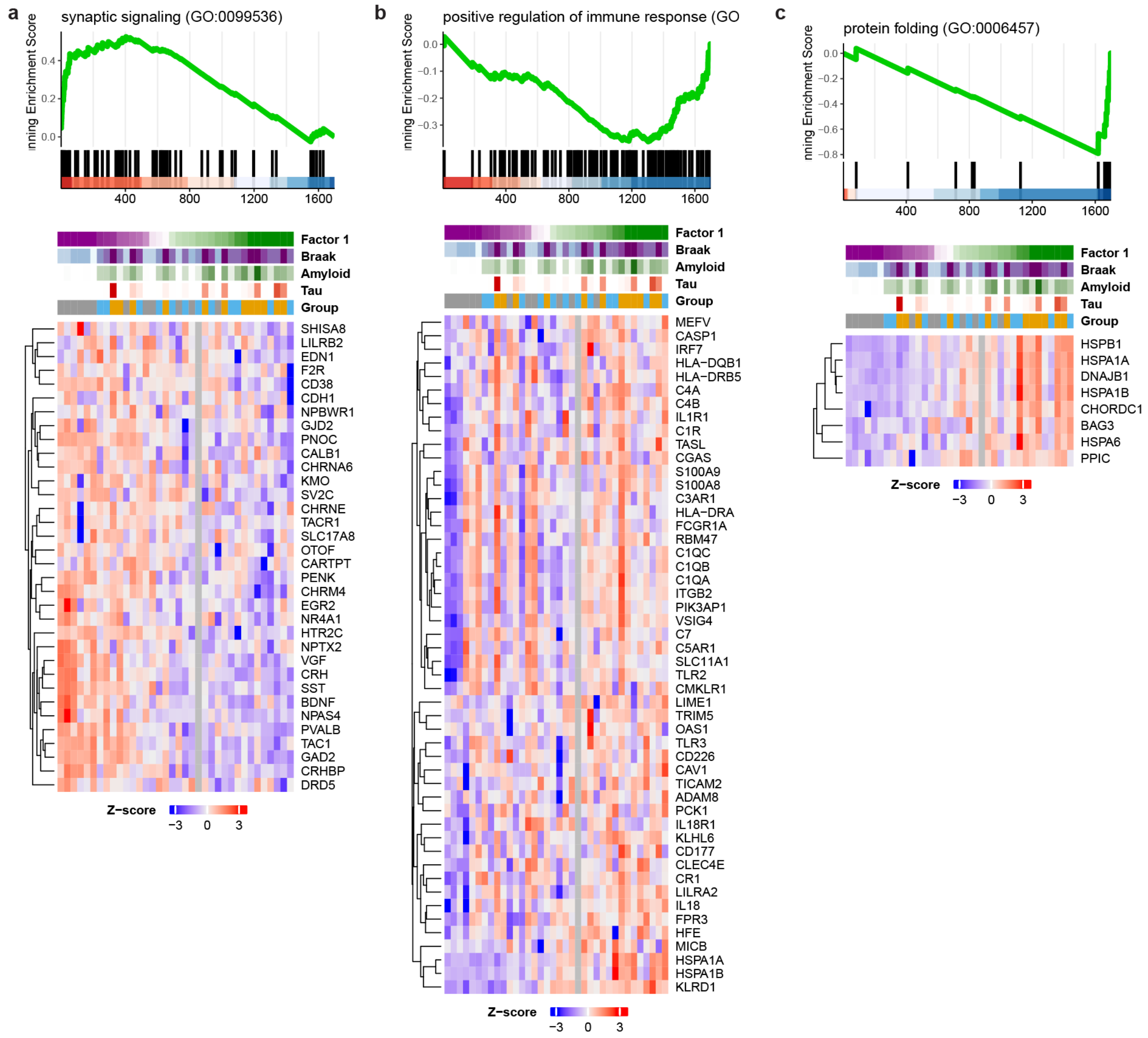


**Supplementary figure 7.** MOFA factor 1 is associated with synaptic signaling (a), immune response (b) and protein folding (c). synaptic signaling is positively associated to Factor 1, meaning downregulation in AD and Resilience, while immune response and protein folding were negatively associated to Factor 1, meaning upregulation in AD and resilience. Data is represented as GSEA plots showing the enrichment, alongside a heatmap of z-scores from the genes in the leading edge of its respective GO term.
